# Single nuclei analyses reveal transcriptional profiles and marker genes for diverse supraspinal populations

**DOI:** 10.1101/2022.07.20.500867

**Authors:** Zachary Beine, Zimei Wang, Pantelis Tsoulfas, Murray G. Blackmore

## Abstract

The mammalian brain contains numerous neurons distributed across forebrain, midbrain, and hindbrain that project axons to the lower spinal cord and work in concert to control movement and achieve homeostasis. Extensive work has mapped the anatomical location of supraspinal cell types and continues to establish specific physiological functions. The patterns of gene expression that typify and distinguish these disparate populations, however, are mostly unknown. Here we combined retrograde labeling of supraspinal cell nuclei with fluorescence activated nuclei sorting and single nuclei RNA sequencing analyses to transcriptionally profile neurons that project axons from the mouse brain to lumbar spinal cord. We identified fourteen transcriptionally distinct cell types and used a combination of established and newly identified marker genes to assign an anatomical location to each. To validate the putative marker genes, we visualized selected transcripts and confirmed selective expression within lumbar-projecting neurons in discrete supraspinal regions. Finally, we illustrate the potential utility of these data by examining the expression of transcription factors that distinguish different supraspinal cell types and by surveying the expression of receptors for growth and guidance cues that may be present in the spinal cord. Collectively these data establish transcriptional differences between anatomically defined supraspinal populations, identify a new set of marker genes of use in future experiments, and provide insight into potential differences in cellular and physiological activity across the supraspinal connectome.

**SIGNIFICANCE STATEMENT:** The brain communicates with the body through a wide variety of neuronal populations with distinct functions and differential sensitivity to damage and disease. We have employed single nuclei RNA sequencing technology to distinguish patterns of gene expression within a diverse set of neurons that project axons from the mouse brain to the lumbar spinal cord. The results reveal transcriptional differences between populations previously defined on the basis of anatomy, provide new marker genes to facilitate rapid identification of cell type in future work, and suggest distinct responsiveness of different supraspinal populations to external growth and guidance cues.

## INTRODUCTION

The supraspinal connectome comprises a diverse and widely distributed set of neurons that project axons to spinal targets and which convey a wide range of motor, autonomic, and sensory modulatory commands. Work spanning more than a century has elucidated the anatomy of supraspinal projections in various model organisms, and in the mouse our recent work has provided a unified description of the supraspinal connectome in three-dimensional space (Wang et al., 2021). In contrast to the state of anatomical and physiological knowledge, much less is understood about the patterns of gene expression that characterize supraspinal neurons or which may distinguish different subtypes. Establishing the transcriptional identities of supraspinal cell types is foundational to understand descending communication from the brain to spinal cord. In addition, the transcriptional signatures of the diverse supraspinal cell types provide essential baseline information to interpret the response of supraspinal populations to injuries and disease states that impact these populations (Blackmore et al., 2021).

Single cell RNA sequencing (scRNA-seq) technologies offer unprecedented insight into cellular diversity by profiling transcript abundance in individual cells and by classifying them based on multi-dimensional indices of similarity (Armand et al., 2021). In the murine nervous system single cell datasets have identified ever finer distinctions between subtypes of neurons withing numerous regions including spinal cord, retina, sensory ganglia, and cortex (Renthal et al., 2020; Russ et al., 2021; Tran et al., 2019; Yao et al., 2021). Apart from recent datasets in corticospinal tract (CST) neurons, however, little is known the transcriptional profiles that characterize neurons that project axons to different spinal targets, and the degree to which anatomical distinctions between the locations of supraspinal cell bodies are associated with distinct patterns of gene expression (Golan et al., 2021; Sahni et al., 2021b).

Here we combined retrograde tracing from lumbar spinal cord axons with single nuclei RNA sequencing (snRNA-seq) to transcriptionally profile diverse types of supraspinal neurons. We identified fourteen discrete classes of supraspinal neurons and identified marker genes that distinguish each class from other supraspinal types. To link these transcriptional clusters to anatomically defined populations we cross-referenced candidate marker genes with anatomically registered transcript expression in the Allen Brain Atlas (https://mouse.brain-map.org/). These correspondences were further validated using in situ hybridization (ISH) detection of marker transcripts in retrogradely labeled supraspinal neurons. Finally, differential gene expression analyses between supraspinal cell types revealed differences in the expression of transcription factors (TFs) and in receptors for growth and guidance cues, hinting at mechanisms that maintain cellular identity and potential differences in responses to extracellular cues between populations. Overall, these data provide new insight into patterns of gene expression that typify and distinguish diverse classes of spinally projecting neurons.

## MATERIALS AND METHODS

### Plasmid construction and cloning

DNA encoding mScarlet (Bindels et al., 2016) was synthesized (Genscript, USA) and fused in frame without a linker to human H2B (H2BC11) (accession #NM_021058) and cloned into the pAAV-CAG-tdTomato (Addgene #59462) using the sites KpnI and EcoRI at the 5 and 3 prime end respectively. AAV2-retro-H2B-mScarlet was made by the University of North Carolina Viral Vector Core with a titer of 4.3×10^12^.

### Spinal cord injections

All animal testing and research was carried out in compliance with ethical regulations laid out by the National Institutes of Health (NIH) guide for the care and use of animals, and all experimental protocols involving animals were approved by the Institutional Animal Care and Use Safety committee at Marquette University (protocol number AR-314). Mice were bred and raised under a 24 h light–dark cycle with 12 h of light and 12 h of darkness. Ambient temperature was maintained at 22 °C ± 2 °C and humidity between 40 and 60%. All 6 mice were female. AAV2-retro particles (1μl) were injected into the spinal cord with a Hamilton syringe driven by a Stoelting QSI pump (catalog #53311) and guided by a micromanipulator (pumping rate: 0.04 μL/min). AAV viral particles were injected L1 spinal cord, 0.35 mm lateral to the midline, to depths of 0.6 and 0.8 mm, with 0.5μl delivered to each site.

### Dissection, FACS sorting, and library preparation

Animals were anesthetized with isoflurane and promptly decapitated. The brain and brainstem were dissected out and placed in ice-cold slushy artificial cerebrospinal fluid (Hearing et al., 2013) for one minute. Brains were then sectioned in the sagittal plane at 500-micron intervals using Adult Mouse Brain Slicer Matrix on ice (Zivic Instruments BSMAS005-2). A total of nine sections were created from each animal (six replicate experimental animals plus four additional for FACS-optimization and other pilot studies), including a midline section and four sections moving lateral into each hemisphere. Retrogradely labelled neurons were then micro dissected using a stereomicroscope and fluorescence adapter (NIGHTSEA SFA-GR). The brain regions collected from each section were recorded for future reference. The three sections from the left hemisphere and one midline section were dissected and flash frozen in a 1.5mL Eppendorf DNA LoBind Microcentrifuge tube. The three sections from the right hemisphere were then collected and flash frozen with identical conditions, as mentioned above. Samples were then stored at -80 °C until FACS sorting.

To prepare cell nuclei on the day of library preparation frozen tissue was promptly transferred from -80 °C storage to a chilled 2mL dounce with chilled Nuclei EZ Lysis Buffer (Sigma-Aldrich N 3408). Sample was dounced 25 times with pestle A and 20 times with pestle B. The dounced sample was then transferred to a chilled 15mL conical tube and an additional 2mL of Nuclei EZ Lysis Buffer was added and gently mixed by inversion. The sample incubated on ice for 5 minutes and then was centrifuged at 500G at 4 °C. The supernatant was removed, and the pellet was resuspended in 4mL of Nuclei EZ Lysis Buffer. The sample again was incubated on ice and centrifuged at 500G at 4 °C for 5 minutes. Following the centrifugation, the supernatant was removed, and the pellet resuspended in 500uL of Nuclei Suspension Buffer (2% BSA, 40U/uL RNase Inhibitor (Invitrogen Ref. AM2684), 1X PBS). The resuspended solution was then filtered through a 20um filter and moved directly to the FACS machine. The collection tube for the sorted nuclei was coated with 5% BSA and contained the 10X Genomics master mix. We initially gate out debris using side scatter area (SSC-A) vs forward scatter area (FSC-A) and forward scatter width (FSC-W) vs forward scatter area (FSC-A). To eliminate doublets, the nuclei that pass through the gates above were then gated by side scatter width (SSC-W) vs side scatter height (SSC-H). Nuclei that passed through all the above gates were then filtered by level of fluorescence so that only the brightest were collected. With the above parameters, the FACS machine was set to collect 5,000 nuclei in five to seven minutes. The collected nuclei were then prepared into libraries, using Chromium Next GEM Single Cell 3’ Reagent Kits v3.1 (PN-1000269), according to the manufacture protocol (10X Genomics, CG000204 Rev D).

### Experimental design and statistical analyses

Libraries were sequenced with an Illumina NovaSeq 6000 at UW-Madison Biotechnology Center and then processed with CellRanger using default parameters to produce a Unique Molecular Identifier (UMI) matrix for all nuclei-containing droplets, which were then analyzed in the Seurate version 4.1.0 R package. Median reads per cell were 125,600, 237,677, and 124,791 and mean unique genes (nFeature) per cell were 3190, 5017, and 4366 for samples 1, 2, and 3 respectively. Nuclei with fewer than 1000 unique genes were excluded, as were nuclei with high levels of Atf3 and Creb5, indicative of cellular stress that likely resulted from the dissection procedure. The three samples were merged using the 2000 most variable genes as input for the “anchor.features” of the FindIntegrationAnchors() function. Clustering was performed according to Seurat recommendations based on 30 principal components as selected by the elbow plot heuristic and using FindNeighbors() and FindClusters() functions. Marker genes for each cluster were identified using FindAllMarkers() with default parameters, which utilizes a Wilcoxon rank-sum test comparing gene expression of cells within a given cluster versus all other cells. To identify variable transcription factors (TFs) we first extracted from the unified dataset a list of all TFs and associated average RNA counts based on TFs as identified in (Lambert et al., 2018). For each TF we then calculated a normalized expression value within each cluster by dividing that cluster’s average RNA count value by the sum of all clusters and multiplying by 100 (i.e. in a hypothetical case in which a gene is expressed at the same level in all fourteen clusters each would receive a normalized score of 100/14 = 7.14.) Variable TFs were defined as scoring less than 1 (under-represented) or greater than 20 (over-represented) in any cluster.

### In situ hybridization and imaging

In situ hybridization was performed using RNAscope Multiplex Fluorescent Detection Kit v2 from ACDbio (Cat no. 323110). Mouse brains were dissected, fresh frozen in OCT (VWR cat no. 95057-838), and cryosectioned at 30um. Cryosection slides were dried at room temperature for one hour before storage at -80°C. Slides were removed from -80°C and promptly began the ISH protocol according to the manufacturer’s protocol. Slides were initially fixed in fresh 4% PFA for one hour at room temperature and washed with 1X PBS (Alfa Aesar, J62036) twice for two minutes each. The slides were then dehydrated in 50%, 70%, and 100% Ethanol for five minutes each and 100% repeated. Slides were allowed to dry at room temperature, followed by addition of hydrogen peroxide for ten minutes. Slides were then washed with nuclease free water, twice for two minutes each. We then applied Protease III (ACDbio cat no. 322337) for eight minutes at room temperature and washed with 1X PBS (Alfa Aesar, J62036) twice for two minutes each. Probes were then applied for two hours at 40°C and washed with 1X wash buffer (ACDbio cat no. 320058) twice for two minutes each. We then performed three consecutive amplification steps, washing with wash buffer in between, as instructed by the manufacturer. The corresponding probe HRP signal was then developed. All probes were detected using a 1:750 dilution of TSA plus fluorescein 488 (Akoya Biosciences NEL741001KT). Slides were then dried overnight. All probes were ordered from ACDbio catalog numbers are provided below. Probes used were Plagl1 (cat no. 462941), Ttc6 (cat no. 1125881), Emx2 (cat no. 319001), Prdm6 (cat no. 456891), Lhx4 (cat no. 1130251), Pard3b (cat no. 832241), Sox14 (cat no. 516411), Kit (cat no. 314151), Col19a1 (cat no. 539701).

Following ISH, IHC was used to amplify the retrograde label. Sections were washed in 1X PBS for 5 minutes and then blocked (10% serum, 1% PBST) for one hour at room temperature. The primary antibody (2-3% serum, 0.4% PBST, 1:500 Rockland 600-401-379) was applied and incubated for ninety minutes at room temperature. Sections were then washed with 1X PBS three times, five minutes each. Secondary antibody (2-3% serum, 0.4% PBST, 1:500 Invitrogen A11035, 1:500 DAPI) was applied and incubated for one hour at room temperature. Sections were then washed with 1X PBS (Alfa Aesar, J62036) three times, five minutes each. Attach coverslip with mounting media (Fluoro-Gel with Tris Buffer Cat# 17985-10).

Slides were imaged within two weeks from completion of ISH/IHC. Slides were imaged with Keyence BZ-X810 microscope and 10x Plan Apochromat Objective (BZ-PA10). Additional images were taken with Nikon AR1+ Confocal Microscope and 60x Apochromat Oil DIC N2 objective (MRD71600, NA 1.4). 60x images were gathered in a single z-plane.

## RESULTS

### Single-nuclei analysis of supraspinal neurons reveals transcriptionally distinct cell types

To label supraspinal neurons, adult mice received lumbar injection of AAV2-retro expressing mScarlet fluorophore that was localized to the nucleus by fused histone protein 2B (AAV2-retro-H2B-mSc) (**Figure 1A**). We have shown previously that this procedure labels tens of thousands of supraspinal neurons distributed through forebrain, midbrain, and hindbrain (Wang et al., 2021, 2018). Two weeks after injection animals were sacrificed, a sagittal series of brain slices were rapidly prepared and placed in ice-cold artificial cerebrospinal fluid and observed under fluorescent light. Based on anatomical location we preliminarily identified fluorescent cells in cortex, hypothalamus, midbrain, dorsal pons, and hindbrain. Regions with fluorescent label were rapidly micro dissected and flash frozen, taking care to exclude as much unlabeled tissue as possible. For each animal, the anatomical location of micro dissected tissue from all sections was recorded. Tissue from two animals were pooled for each sample, from which cell nuclei were extracted and processed by fluorescent activated nuclei sorting (FANS) to purify the supraspinal subset. In initial studies nuclei were sorted into PBS and inspected visually to confirm mScarlet signal in more than 98% (**Figure 1C**). In subsequent experiments nuclei were sorted directly into RT Master Mix followed immediately by GEM formation and library preparation according to 10X chromium instructions. Pilot studies revealed that library quality was highly dependent on rapid sorting, such that run times above 10 minutes led to unacceptable levels of ambient RNA in the final sample. Thus, to minimize run times it was essential to initiate FANS sorting with micro-dissected samples highly enriched for labeled nuclei. Five thousand nuclei were gathered from each sample and of these, based on the volumes transferred to GEM formation, an estimated 3400 nuclei entered initial library creation. This procedure was repeated three times, yielding three independent replicates, each derived from two pooled animals. Libraries were sequenced to a minimum depth of 100 million reads, followed by clustering and analysis by Seurat (Butler et al., 2018).

**Figure 1.**
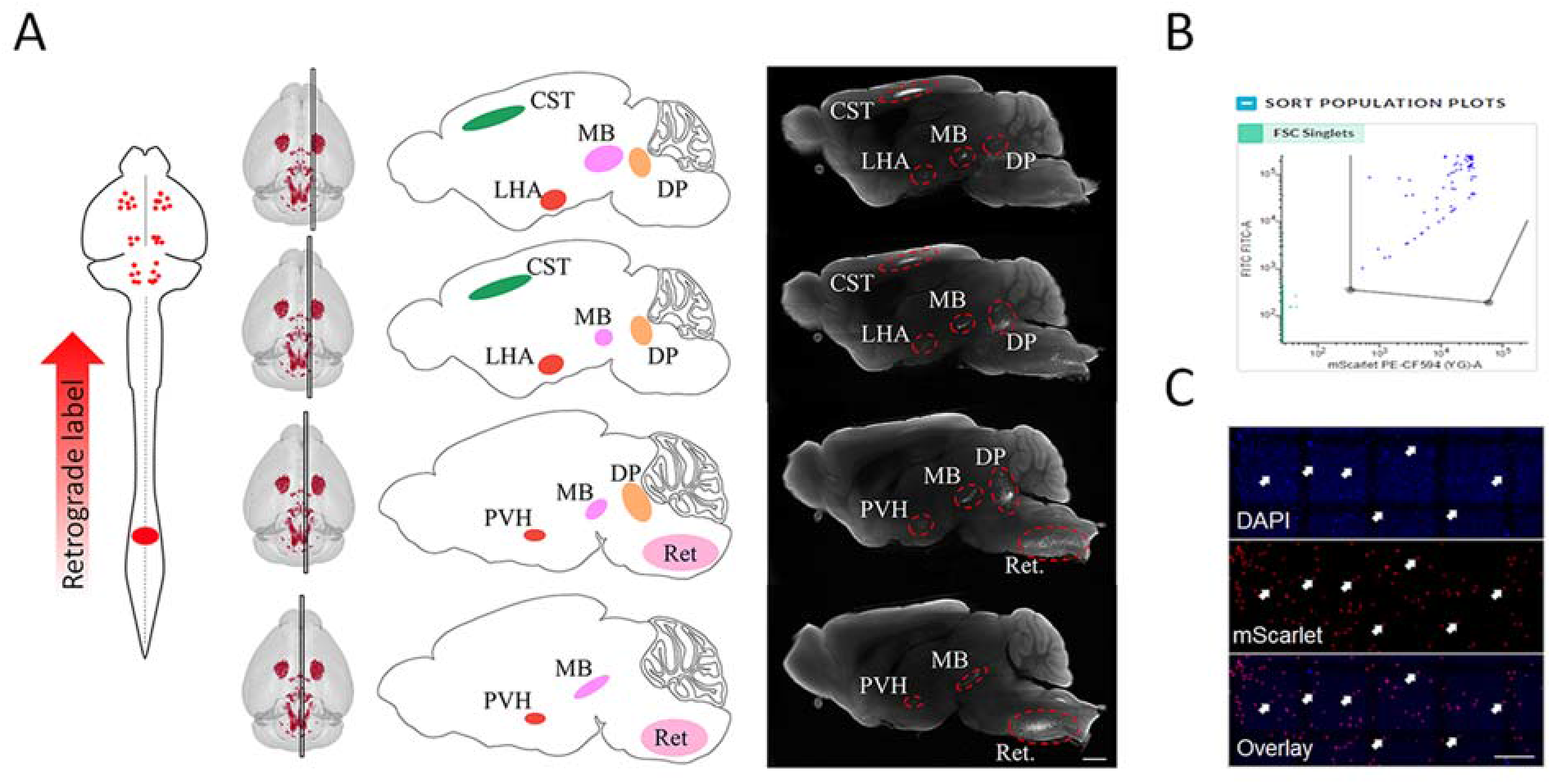
Retrograde labeling and FANS purification of supraspinal nuclei. A) AAV2-rctro-H2B-mScarlet was injected to lumbar spinal cord, followed two weeks later by microdissection of fluorescently labeled brain regions in sagittal section. Middle panels show schematics indicating the anatomical regions of interest and right panel shows tissue slices with retrograde fluorescent label and red lines indicating regions of microdissection. B) Representative image of gates used for isolation of mScarlet-labeled nuclei. C) Visual inspection of nuclei after FANS shows mScarlct label in nearly all nuclei. D) Indicates the brain regions that were microdisscctcd in each sample. CST = corticospinal tract. LHA = Lateral Hypothalamic Area PVH = Paraventricular Hypothalamus. MB = Midbrain. DP = Dorsal Pons. Ret = Medullary Reticular Formation. Scale bars arc 1mm (a) and 50 μpm (c).

In initial quality control steps, clusters of nuclei were identified with discrepantly low feature counts and with high levels of stress-related transcripts such as Atf3 (Hai et al., 1999). These ∼1000 nuclei were presumed to be responding to damage or loss of nuclear membrane integrity during sample preparation and were removed from subsequent analysis. The remaining nuclei, which numbered 7,748 across the three samples, were reclustered to yield 18 initial groups (**Figure 2A**). Clusters were highly consistent between the three samples, an important indication that all samples contained a consistent set of putative cell types and that no cluster was the result of artifacts that were specific to any one sample (**Figure 2B**). Based on the method of long-distance retrograde labeling, we predicted that sorted nuclei should derive from long distance projection neurons and not from non-neuronal cell types. Indeed, all clusters expressed high levels of neuronal markers Syt1 and Rbfox3/NeuN (Duan et al., 2015; Park and Ryu, 2018) and at most trace amounts of various markers from non-neural cells including oligodendrocytes, astrocytes, microglia, and ependymal cells (Chen et al., 2017; Eng and Ghirnikar, 1994; Galland et al., 2019; Konishi et al., 2017; Marques et al., 2016; Sock and Wegner, 2021) (**Figure 2C**). We examined markers for neurotransmitter subtypes including glutamatergic (either Vglut1/Slc17a7 or Vglut2/Slc17a6) (Herzog et al., 2001; Kaneko and Fujiyama, 2002), inhibitory (Gad2) (Erlander et al., 1991), serotonergic (Tph2) (Hendricks et al., 1999; Ren et al., 2019), or noradrenergic neurons (Slc2a6) (Mulvey et al., 2018), all of which are known to project from brain to lumbar spinal cord. All 18 clusters were marked by expression of one of these transmitters, and none displayed expression of multiple transmitters (**Figure 2C**,**D**). Interestingly, nearly all clusters expressed either Slc17a6 or -7, while only three very small clusters expressed the inhibitory, serotonergic, or noradrenergic markers: these non-glutamatergic clusters comprised less than 3% of the analyzed nuclei (**Figure 2D, E**) This level of glutamatergic enrichment is likely disproportionate to brain-lumbar projection types, as inhibitory neurons may comprise more than one third of brainstem-spinal projection neurons (Holstege, 1991). One likely explanation is that the tropism of AAV2-retro favors glutamatergic neurons over others (Ganley et al., 2021; Tervo et al., 2016; Wang et al., 2018). In summary, retrograde viral labeling and FANS of produces a high purity neuron population, predominantly glutamatergic, which readily segregates into transcriptionally distinct clusters.

**Figure 2.**
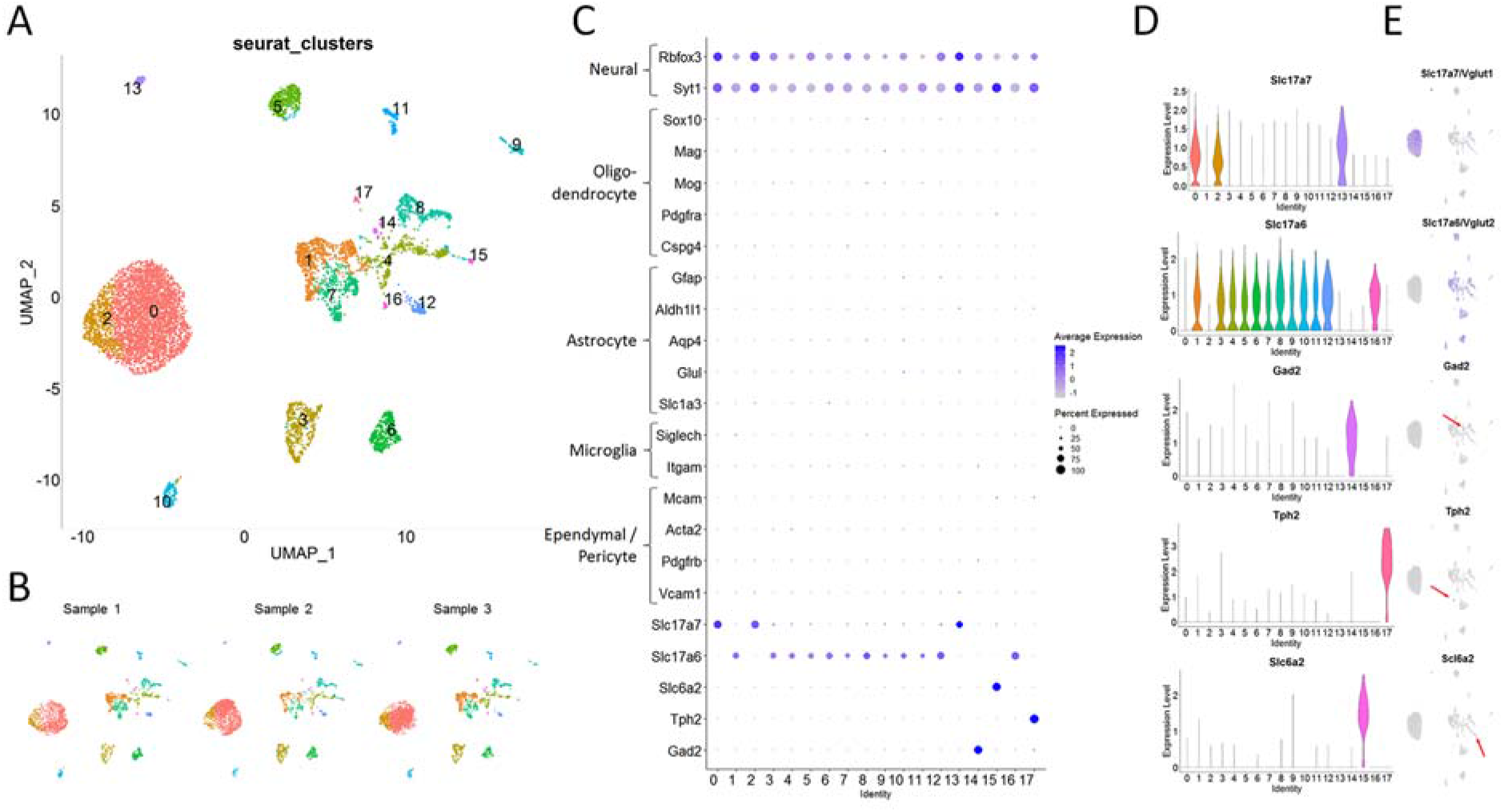
snRNA-seq analysis indicates that samples are mostly glutamatergic neurons. A) UMAP clustering of 7748 supraspinal nuclei identifies 18 distinct groups. B) Separation of the three merged samples confirms tliat each contributes to all 18 clusters, supporting replicate consistency. C) A dotplot sliows expression of neuronal markers in all clusters and very low expression of non-neuronal markers, as expected from the retrograde labeling strategy. D, E) Violin plots (D) and feature plots (E) indicate expression of glutamatergic markers in most clusters, and expression of gabaergic. serotonergic, or noradrenergic markers in only small clusters.

As a first step in classifying supraspinal populations we sought to identify CST neurons, a prominent and relatively well-characterized projection from layer V of cortex to lumbar spinal cord. Indeed, a large and distinctly positioned set of cells displayed high expression of layer V markers genes Bcl11b, Crym, and Fezf2 (Arlotta et al., 2005; Fink et al., 2015; Greig et al., 2013) (**Figure 3A-C**). Examination of transcripts that were enriched in this group revealed additional transcripts localized to layer V cortex according to *in situ* data from the Allen Brain Atlas, including Bcl6, Pdlim1, Cacna1h, Mylip, and Kcng1 (**Figure 3G-K**). We conclude that Seurat clusters 0 and 2, which together contain 3823 nuclei (49.9% of the total gathered), correspond to CST neurons. Another small cluster contained 88 nuclei and displayed high expression of general markers for cortex (Slc17a7, Satb2, Camk2a) but low levels of layer V markers and high levels of Slc30a3, a recently identified marker of intrahemispheric cortical neurons (Zhang et al., 2021). This cluster likely reflects trace contamination (∼2.2%) by non-CST neurons in the cortical sample and was removed.

**Figure 3.**
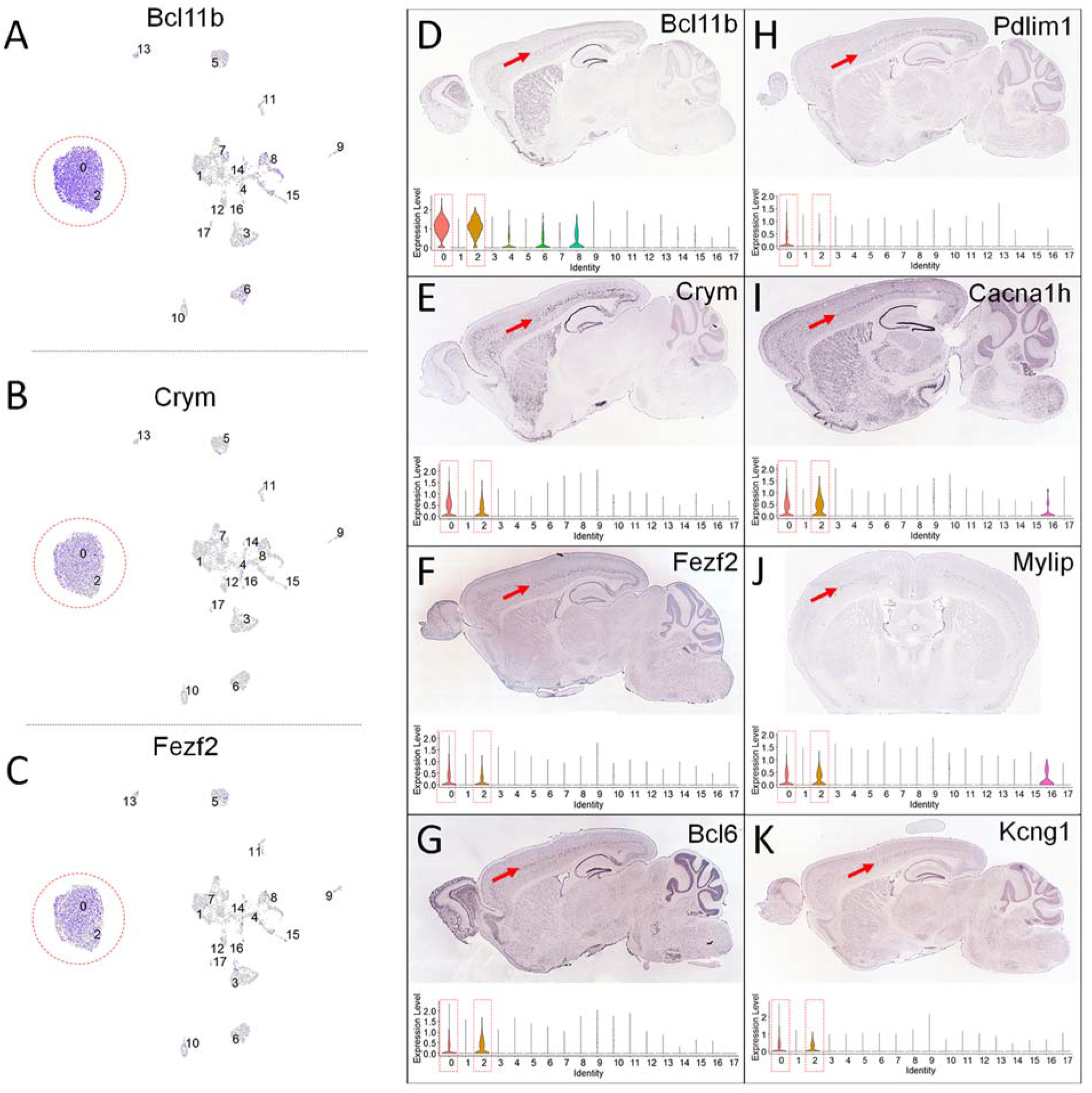
Corticospinal neurons identified by established layer V markers. A-C) Feature plots of Bel 11b (A), Crym (B), and Fezf2 (C) show high enrichment in clusters 0 and 2 (red circles). D-K. show ISH detection from the Allen Brain Atlas of transcripts w ith enrichment in layer V of cortex (red arrows). Corresponding violin plots indicate high expression of transcripts in clusters 0 and 2 (dotted red rectangles), consistent with corticospinal identity.

We next aimed to identity the remaining fifteen Seurat clusters, which presumably corresponded to various supraspinal populations of subcortical origin, most of which lack established markers. Starting from transcripts that were highly enriched in each cluster, we adopted a strategy of manual curation that compared ISH data from the Allen Brain Atlas to the known locations of supraspinal neurons. This approach was aided by our recently created 3D atlas of supraspinal populations (Wang et al., 2021), which registers retrogradely labeled cell nuclei to a digital neuroanatomical atlas based on the Allen Brain Atlas. In this way we could systemically examine locations in Allen Brain Atlas images that we knew to harbor supraspinal neurons, in search of putative marker genes from the scRNA-seq data. Below we outline the results of this initial classification in approximate rostral-to-caudal sequence.

Two clusters likely derived from distinct regions of the hypothalamus, a known source of supra-lumbar input (Hosoya, 1980). The first expressed high levels of Sim1, a known marker for the paraventricular hypothalamic (PVH) region (Michaud et al., 1998) (**Figure 4A-D**). The second expressed high levels of Plagl1, which in Allen Brain data is expressed strongly in the lateral hypothalamus (LH) (**Figure 4E-H**). Next, a prominent cluster selectively expressed Ttc6, which we found to be highly expressed in the red nucleus in Atlas images (**Figure 4I-L**). Cartpt is a known marker for the Edinger Westphal nucleus (EW) (Topilko et al., 2022; Xu et al., 2014), and was selectively expressed in a small cluster (**Figure 4M-P**). Another small cluster expressed Slc6a2, a well-established marker for noradrenergic neurons, and thus likely corresponds to the locus coeruleus (Mulvey et al., 2018) (**Figure 4Q-T**). As noted previously, it is possible that the small size of this cluster may not reflect the true abundance of this projection, but rather the predominantly glutamatergic tropism of AAV2-retro. Finally, a larger group of cells selectively expressed the TF Prdm6, a transcript that localized in the Allen Brain Atlas to the dorsal pontine area (**Figure 4U-X**). Our prior registration data showed a prominent cluster of supra-lumbar neurons in this region which are distributed across several adjacent sub-nuclei such as Barrington’s nucleus. Consistent with this, the current data showed expression of Crh, a known marker for Barrington’s nucleus (Verstegen et al., 2017), in a subregion of the Prdm6 cluster. Accordingly, we preliminarily assigned the Prdm6 cluster, which likely contains Barrington’s nucleus among others, to the dorsal pontine region.

**Figure 4.**
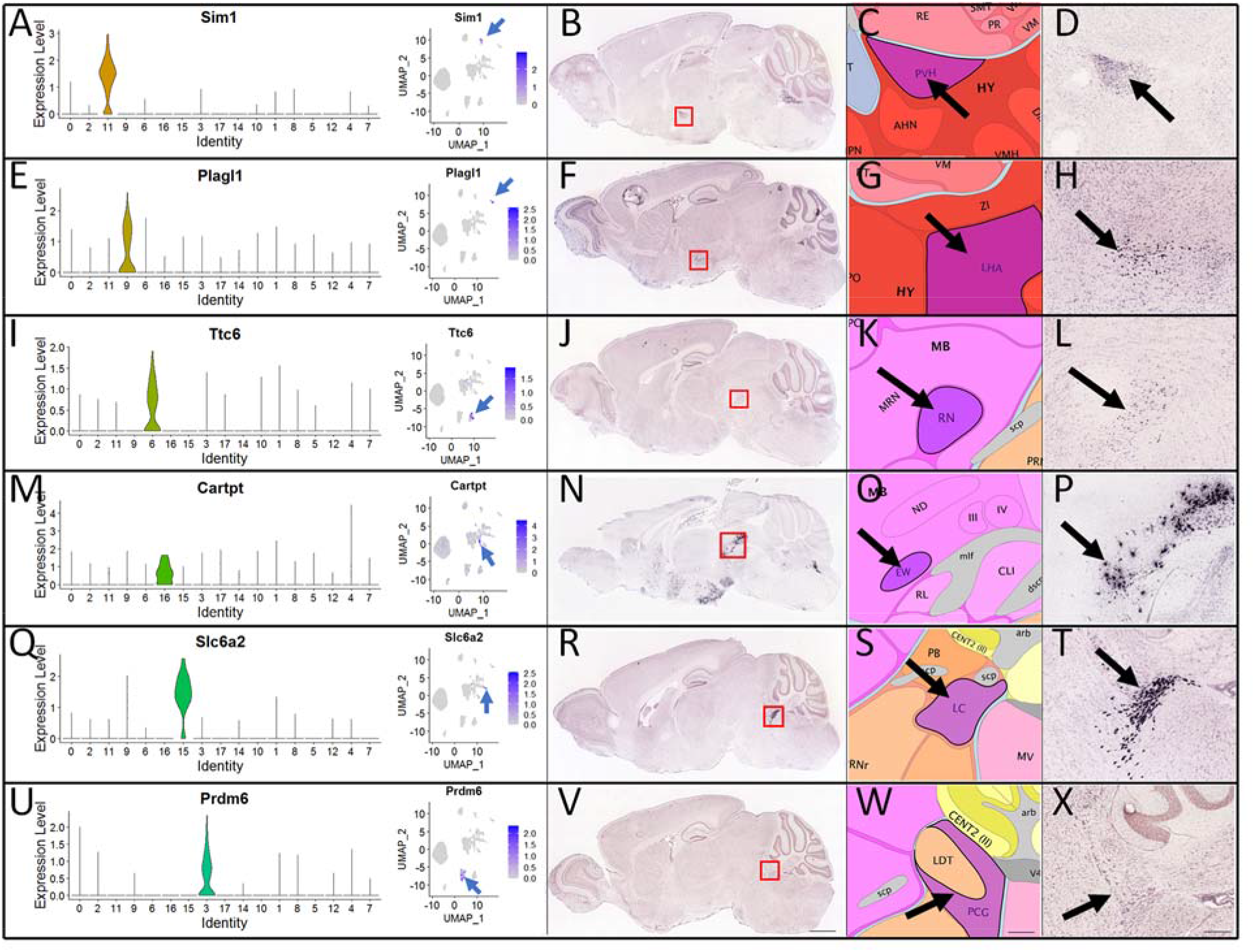
In situ hybridization data from the Allen Brain Atlas enables preliminary identification of transcriptional clusters. A, E, I, M, Q, U show violin plots (left) and feature plots (right) of transcripts with high enrichment in single clusters. B, F, J, N, R, V show ISH data from the Allen Brain Atlas, with red boxes indicating regions of interest. C, G, K, O, S, W show the corresponding anatomical registration to regions that are known to supply supraspinal input. D, H, L, P, T, X show higher magnification views with detection of putative marker genes. Arrows point to areas of specific gene expression: LHA = Lateral Hypothalamic Area PVH = Paraventricular Hvpothalamus. RN = Red nucleus. EW = Edinger-Westphal nucleus. LC = Locus Coeruleous, PCG - Pontine Central Grey. Formation Sente bar in V-1mm Sente bar in W and X – 200 μm

The remaining nine subcortical clusters, which comprised about 34% of all cells, were marked by expression of Hox3 and Hox4 gene clusters (Krumlauf and Wilkinson, 2021) (**Figure 5A, B**). The Hox genes are important for patterning and segmentation of rhombomeres that give rise to the medulla and pons and serve as canonical markers of brainstem neurons (Chambers et al., 2009). Interestingly the specific Hox gene clusters (**Figure 5A**) are mostly expressed in the caudal part of the developing hindbrain suggesting that these clusters are anatomically located in the medulla. Consistent with this, most of the Hox+ clusters also expressed the homeobox protein Lhx4, previously noted for expression in reticulospinal neurons of brainstem origin (Bretzner and Brownstone, 2013; Cepeda-Nieto et al., 2005) (**Figure 5C**). Also prominent in the Hox+ clusters was another homeobox gene, Vsx2 / Chx10, previously detected in some populations of brainstem-spinal neurons (**Figure 5D**) (Usseglio et al., 2020). We therefore concluded that these nine clusters likely derived from the caudal part of the brainstem. The Hox+ group also contained two small clusters, noted above, that expressed markers for inhibitory or serotonergic subtypes which likely correspond to trace amounts of brainstem gabaergic and raphe-spinal populations that took up AAV2-retro. In general, the remaining cell clusters that expressed Hox genes were not as sharply segregated as other supraspinal populations. Nevrtheless, some transcripts were found to display clear concentration in subregions of the putative brainstem populations. These included Daam2 (**Figure 5E**), Pard3b (**Figure 5F**), Sox14 (**Figure 5G**), Col19a1 (**Figure 5H**), and Kit (**Figure 5I**). Examination of data from the Allen Brain Atlas supported expression of these markers in brainstem regions, but unlike clusters assigned to the midbrain or forebrain did not reveal strong segregation to annotated subregions of brainstem.

**Figure 5.**
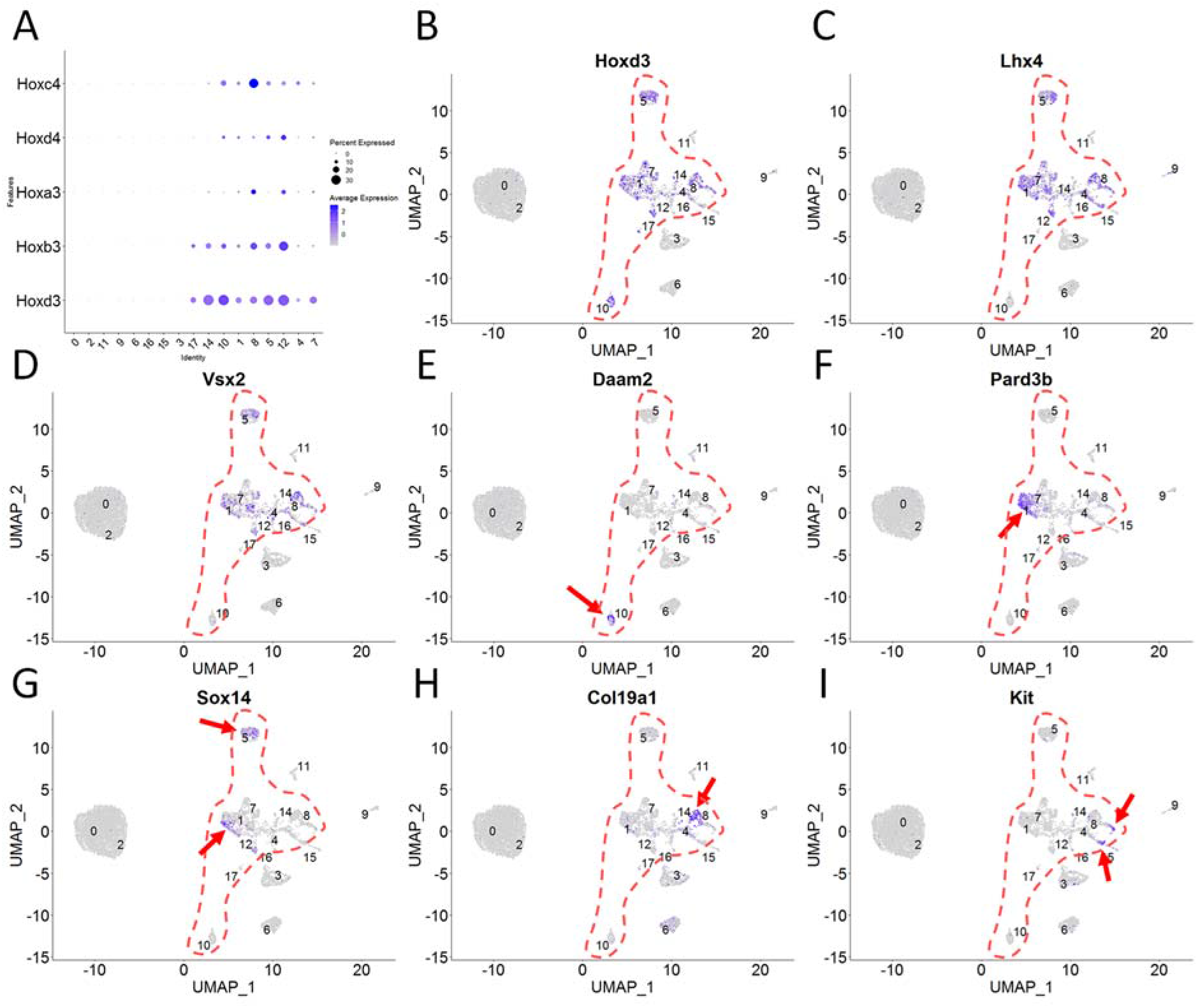
Identification of hindbrain populations. (A) A dotplot of Hox transcript levels indicates selective expression in a subset of HOX gene clusters, indicating hindbrain populations of caudal rhoinboineric origin. (B-H) show feature plots with putative hindbrain populations outlined and specific transcripts enriched in subregions indicated by red arrows.

Figure 6 summarizes assignments of identity to fourteen cell populations, some of which are aggregates of original Seurat clusters, and lists “marker” transcripts that were concentrated in each. Supraspinal populations rostral to the hindbrain appear to be transcriptionally distinct, as evidenced by widely separated clustering via UMAP (**Figure 6A**) and the presence of multiple transcripts in each that are highly specific (**Figure 6B**). In contrast, distinctions between hindbrain populations are generally less definitive, with less separation by UMAP and fewer distinctive marker genes. As such the hindbrain was divided into five populations, the first four of which were based on well segregated markers (Daam2, Pard3b, Col19a1, and Sox14), and the fifth derived from a disparate collection of clusters, none of which displayed highly distinctive markers.

**Figure 6.**
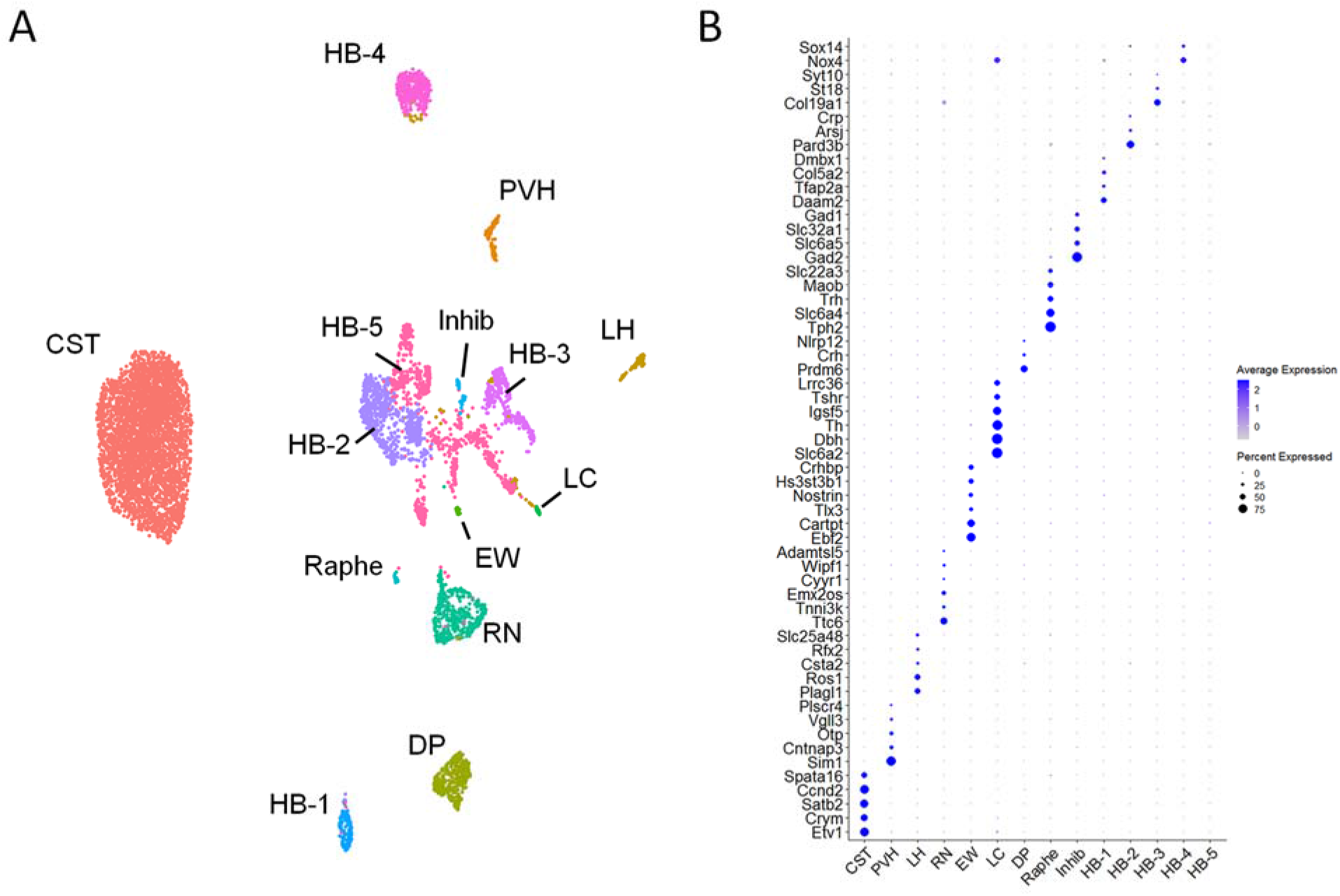
Anatomical assignment and marker genes for populations of supraspinal neurons. (A) UMAP clustering of supraspinal nuclei with anatomical designations for each. (B) A dotplot of gene expression showing examples of transcripts with enriched expression in each cluster relative to other supraspinal populations. CST = Cortico spinal tract. HB = Hindbrain. LH = Lateral Hypothalamus PVH = Paraventricular Hypothalamus. RN = Red nucleus. EW = Edinger-Westphal nucleus. LC = Locus Coeruleous. DP = Dorsal Pons.

Overall, single nuclei profiling identified a total of thirteen types of supraspinal neurons that are readily distinguishable, along with an additional set of neurons that derive from hindbrain regions, but which were not clearly distinguishable in the current dataset.

#### ISH verifies expression of putative marker genes in supraspinal populations

To validate the classification of populations outlined above, we combined retrograde labeling of lumbar projecting supraspinal neurons with visualization of transcripts using ISH. As previously, adult mice received injection of AAV2-retro-H2B-mScarlet to lumbar spinal cord followed two weeks later by ISH of brain sections. ISH was performed using ACDbio probes for specific marker genes, followed by imaging to assess co-localization of mScarlet+ supraspinal nuclei with *in situ* signal. This analysis found strong concordance of the pattern of transcript detection with predictions from the single nuclei data. For example, Plagl1 transcript was detected in supraspinal neurons located in the lateral hypothalamus (**Figure 7A-C**). Two candidate markers for the red nucleus, Ttc6 and Emx2 were both detected in rubrospinal neurons (**Figure 7 D-I**), while Prdm6 was detected as predicted in supraspinal neurons clustered in the dorsal pons (**Figure 7J-L**). Finally, in hindbrain supraspinal neurons we detected frequent expression of Lhx4, and in subsets of neurons detected Pard3b, Sox14, Kit, and Col19a1 (**Figure 8A-O**). It should be noted that for all the probes tested we also observed positive signal in brain regions outside of the supraspinal regions of interest. Thus, when considered in the broader context of the nervous system, these transcripts cannot be considered specific to a particular supraspinal population. Critically, however, mScarlet+ nuclei outside the regions of interest revealed no detection of the marker transcript (**Extended Data 7-1, Extended Data 8-1**). Thus, within the subset of neurons that project axons from the brain to lumbar spinal cord, these transcripts serve as markers for distinct subtypes of neurons with defined anatomical locations. Overall, the ISH results validate the anatomical assignments made previously to the cell type grouping presented in Figure 6.

**Figure 7.**
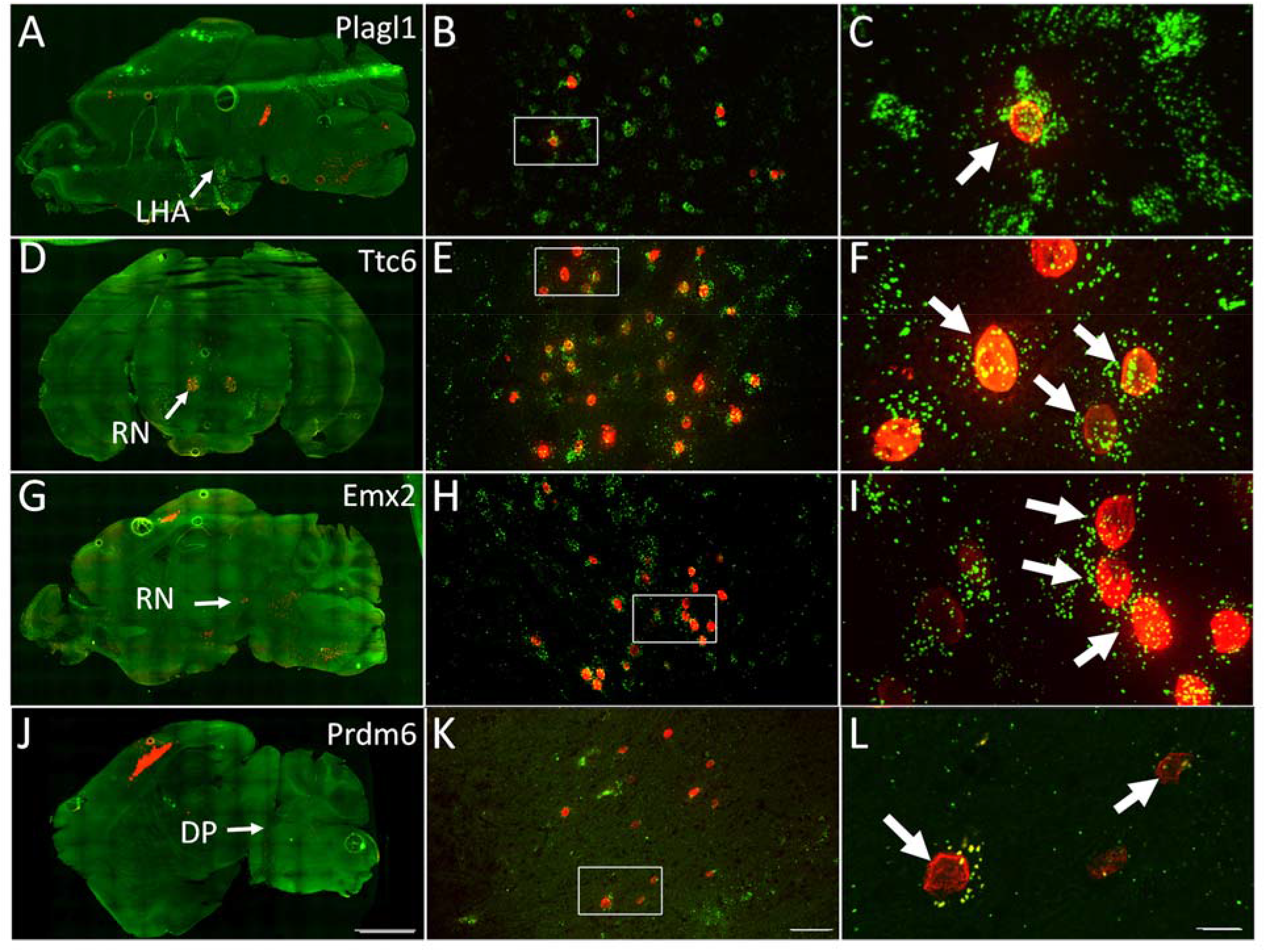
Visualization of candidate marker genes in subcortical supraspinal populations. Adult mice received lumbar injection AAV2-retro-H2B-mScarlet followed two weeks later by tissue sectioning and ISH detection of candidate marker genes. A, D, G, and J show brain sections with target populations indicated by arrows. B, E, H and K show higher magnification view of target populations and C, F, I, and L show high resolution of supraspinal cell nuclei (red) with transcript detection (green). Plagll (A-C) is detected in supraspinal neurons in the lateral hypothalamic area. Ttc6 (D-F) and Emx2 (G-I) in the red nucleus, and Prdin6 in the dorsal region of tire pons. Scale bars are (J) 2mm. (K) 50 μpm. and (L) 10pm. Extended Data 7-1 provides images of off-target supraspinal populations that lack marker expression, indicating specificity.

**Figure 8.**
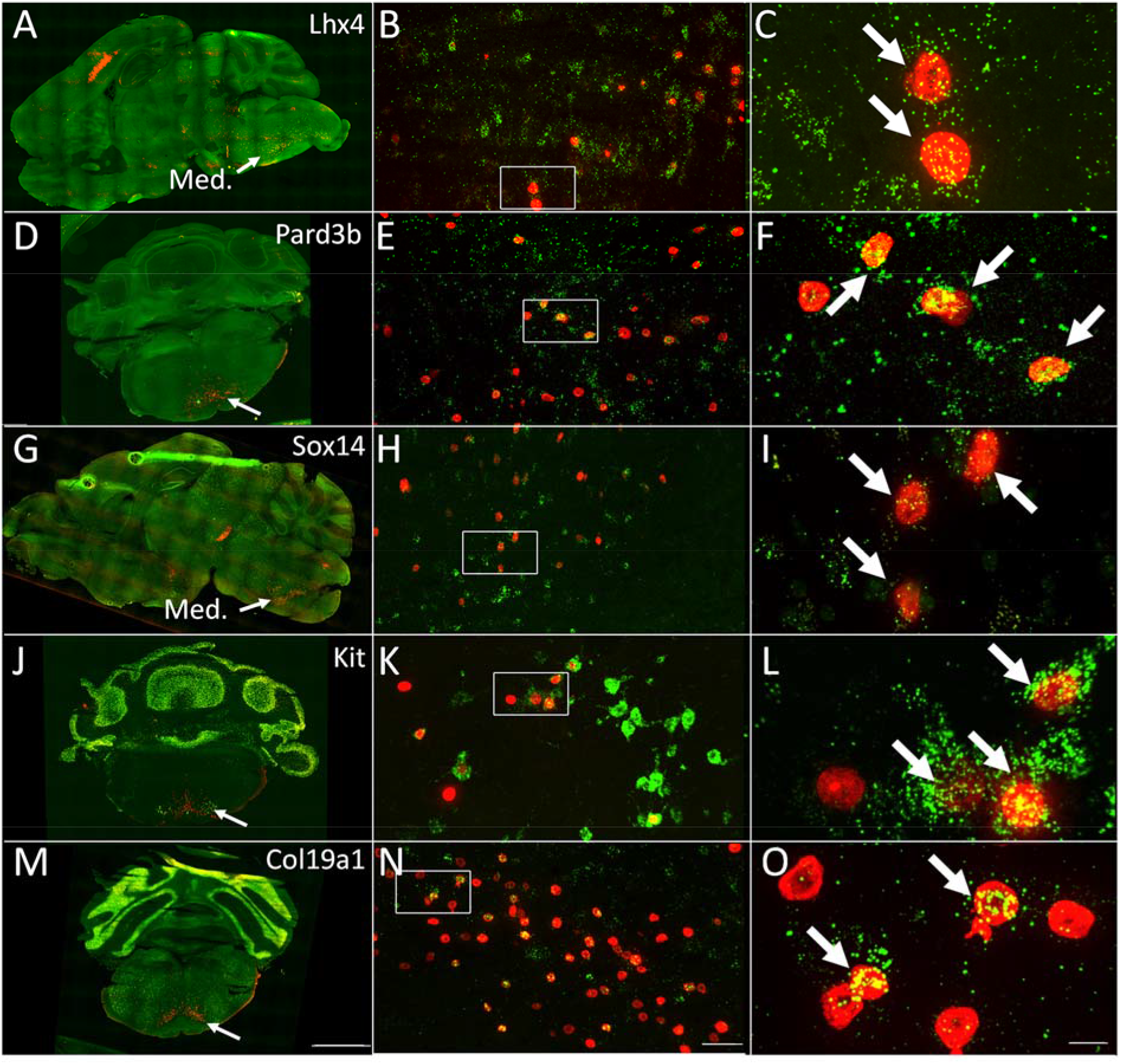
Visualization of candidate marker genes in hindbrain supraspinal populations. Adult mice received lumbar injection AAV2-retro-H2B-mScarlet followed two weeks later by tissue sectioning and ISH detection of candidate marker genes. A, D, G, J, and M show overview of brain sections with medullary regions of interest (arrows). B, E, H, K and N show higher magnification views of target regions. C, F, I, L and 0 show high resolution images of supraspinal cell nuclei (red) with transcript detection (green). Lhx4 (A-C) is broadly expressed and Pard3b (D-F). Sox 14 (G-I). Kit (J-L) and Coll9al (M-O) are detected in discrete subsets of supraspinal neurons. Scale bars are (M) 2mm, (N) 50 μpm. and (0) 10pm. Extended Data 8-1 provides images of off-target supraspinal populations that lack marker expression, indicating specificity.

### Transcription factor expression and receptor/ligand profiles of supraspinal populations

Diverse families of transcription factors are essential for neuronal identity and connectivity during development and postnatal periods (Perreault and Giorgi, 2019; Russ and Kaltschmidt, 2014). To gain insight into the regulation and maintenance of different supraspinal subtypes we first queried the data for differences in the expression of transcription factors (TFs). Starting from a curated list of all TFs (Lambert et al., 2018), we focused on TFs that displayed the greatest variability in expression between different supraspinal populations (see methods). **Figure 9A** shows a set of 79 variable TFs, while Extended Data 9-1 provides a spreadsheet with the average expression of all transcripts, including those identified as TFs, across all supraspinal cell types. As already noted, some transcription factors are highly specific to a single supraspinal type, such as Sim1 in PVH, Prdm6 in Dorsal Pons, Plagl1 in lateral hypothalamus, and Sox14 in a subset of hindbrain neurons. Additional TFs with high specificity included Ebf2 in the midbrain midline nuclei (EW), Phox2a and -b in the Locus Coeruleus, Rfx4 in LH, and Rreb1 in the Red Nucleus. Numerous TFs differed broadly between CST and subcortical populations, with some factors highly enriched in cortex (Etv1, L3mbtl4, Satb2, and Zeb2) and others expressed in numerous subcortical populations but low or absent in CST (Cux2, Dach1, Glis1, Zfhx4, and Zbtb20). Other TFs were not specific to any one population but instead were common to small subset and excluded from all others. One striking example of this is Tfap2b, which was found to be abundant in EW, LC, and a subtype of hindbrain but was not expressed elsewhere. Overall, these data provide foundational knowledge regarding the overlapping sets of transcription factors that typify single types and groups of supraspinal neurons.

**Figure 9.**
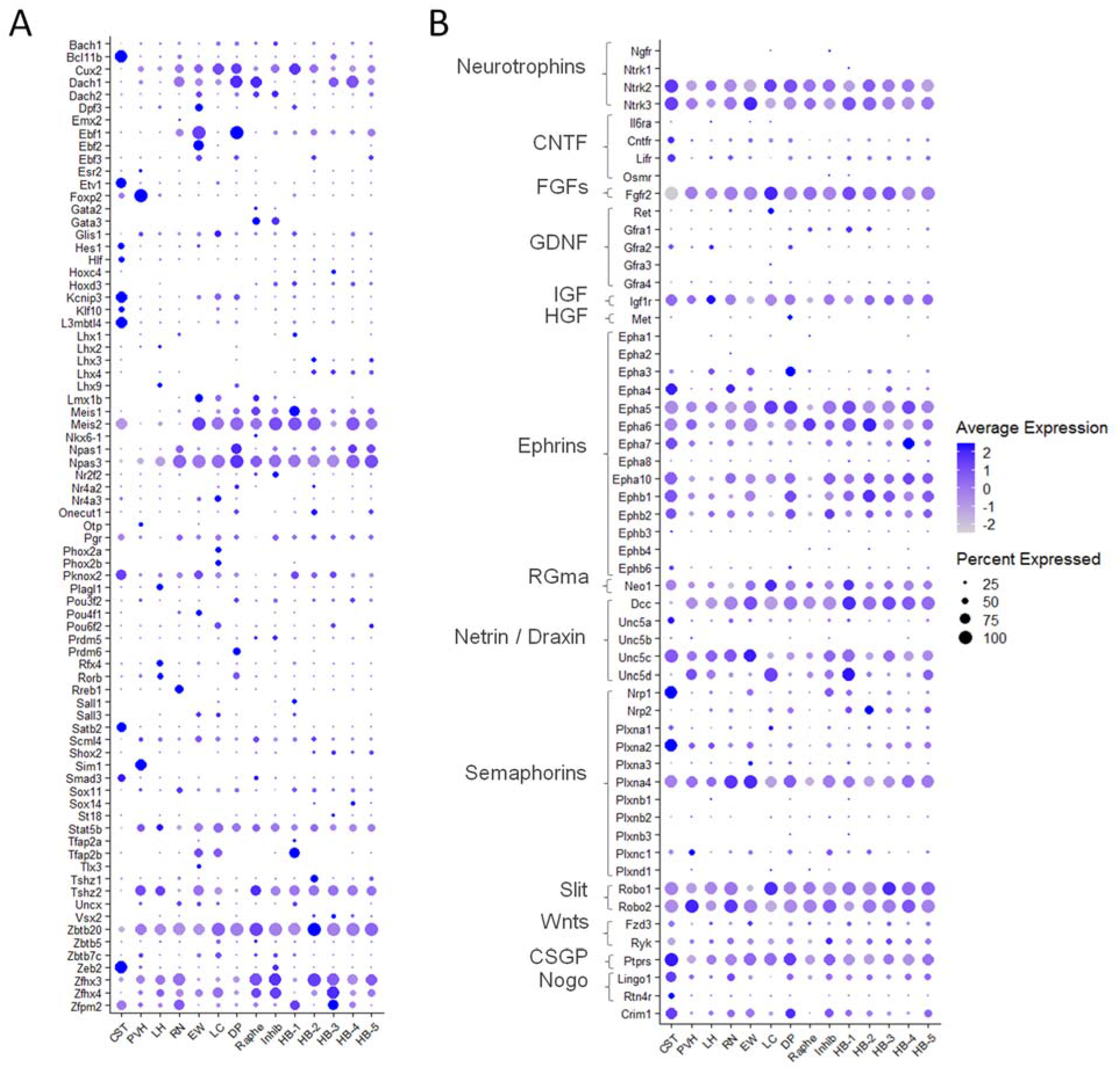
Differential expression of transcription factors, growth factor, axon guidance and growth inhibitory receptors in supraspinal populations. (A) A dotplot indicates the expression levels of selected transcription factors in each of fourteen distinct ripes of supraspinal neurons. (B) A dotplot indicates the level of expression of selected receptors in supraspinal populations. Relevant ligands are listed to the left and include neurotrophins. axon guidance cues, and growth-inhibitory cues. Extended Data 9-1 provides average expression of all transcripts in all populations.

We next considered the expression of receptors for growth factors and axonal guidance cues across supraspinal subtypes. This information is potentially informative in predicting differential responses to cues in the spinal cord environment. We first assembled a list of ligands of interest, based on neurotrophins, growth factors, and axon guidance cues that are either endogenous to spinal cord tissue or which have been exogenously applied in experimental paradigms of spinal injury (Badhiwala et al., 2018; Dun and Parkinson, 2017; Gao et al., 2019; Giger et al., 2010; Goldshmit et al., 2011; Haenzi and Moon, 2017; Hata et al., 2006; Kitamura et al., 2019; Liu et al., 2017; Rosich et al., 2017; Walker and Xu, 2018; Yamane et al., 2019). We then examined the expression of various receptors for each cue across different subtypes of supraspinal neurons (**Figure 9B**). Many receptors displayed broad expression across all types of supraspinal populations (e.g. neurotrophin receptors Ntrk2/Trkb and Ntrk3/TrkC), and others were missing from all types (e.g. Ntrk1/TrkA). It is notable that receptors for GDNF, which was shown recently to stimulate growth from injured propriospinal neurons, are mostly absent from supraspinal populations in the brain, which is consistent with minimal levels of supraspinal regeneration that were reported (Anderson et al., 2018). Expression of some receptors was highly variable, for example CST neurons expressed high levels of receptors for semaphorins, Npn1 and Plexa2, and high levels of Unc5d, which confers repulsive guidance signaling from netrin (Alto and Terman, 2017; Dun and Parkinson, 2017; Helmbrecht et al., 2015; Huber et al., 2005; Huettl et al., 2011). In contrast, the Dcc receptor that confers positive growth responses to netrin (Dun and Parkinson, 2017) as virtually absent from CST neurons but expressed broadly across other supraspinal types. In addition, some interesting differences between populations were detected. These data are consistent with the established responsiveness of CST neurons to semphorin signaling and hint that CST neurons may potentially respond differently to netrin ligands than do other supraspinal populations. Important caveats to this approach include the fact that mRNA abundance does not necessarily scale with protein expression and that some receptors are known to be expressed by neurons but not transported into axons (Koseki et al., 2017). Thus, the presence of mRNA for a receptor is not definitive evidence of that the neuron’s axon will respond to the corresponding ligand when presented in spinal tissue. Nevertheless, particularly in cases in which the mRNA for a receptor is not detected in a given supraspinal class, these comparative data provide the basis to rapidly generate hypotheses of differential response to ligands presented in the spinal cord in regeneration studies.

## DISCUSSION

We have identified fourteen transcriptionally distinct populations of neurons that project axons from the murine brain to the lumbar spinal cord. In addition, using a combination of well-established and newly discovered marker genes we have associated these transcriptionally distinct clusters with anatomically defined populations, thus linking new transcriptional insights to a rich neuroanatomical literature. These data establish new marker genes, offer insight into potential physiological differences between supraspinal cell types, and provide a population-by-population baseline for future study of the transcriptional impacts of injury and disease.

### Neuronal diversity in the supraspinal connectome

Single cell RNA sequencing technologies are rapidly expanding awareness of cellular diversity in the nervous system. Single-cell and -nuclei data exist for various regions in the adult and developing mouse nervous system including retina (Tran et al., 2019), dorsal root ganglion (Renthal et al., 2020), spinal cord (Russ et al., 2021), cortex (la Manno et al., 2021; Yao et al., 2021), and CST (Golan et al., 2021). Here we aimed to broaden understanding of supraspinal cell types spanning forebrain, midbrain, and hindbrain, and to test whether transcriptionally unique profiles associate with established anatomical classifications. Because supraspinal populations carry distinct functions and can target distinct spinal circuits (e.g. motor versus autonomic), we expected to identify differences in gene expression. Indeed, our findings broadly confirm the assumption that anatomically separated supraspinal neurons also differ from one another in their expressed transcripts. Many populations existed as discrete UMAP clusters with some transcripts that were ubiquitous within that cluster but essentially absent elsewhere. ISH-based visualization further validated selective expression for many transcripts.

Brainstem-spinal populations, however, stand as a partial exception. Although brainstem neurons clustered separately from midbrain and forebrain and were recognizable by markers such as Hox and Lhx factors, brainstem subtypes remained closely adjacent by UMAP. We did identify transcripts including Sox14, Pard3b, Col19a1, and Kit that were detected in only some brainstem clusters and which by ISH were expressed selectively in brainstem discrete regions. In general, however, hindbrain clusters shared putative marker genes with at least one other cluster. These results are reminiscent of the ventral spinal cord, in which neuronal subtypes were not well delineated in early single cell datasets (Sathyamurthy et al., 2018) but could later be distinguished in larger samples (Russ et al., 2021). Finer distinctions between hindbrain populations will likely emerge as additional datasets are created and aggregated.

Nevertheless, the present data confirm that multiple subtypes exist within the umbrella classification of glutamatergic brainstem-spinal projection neurons and identify multiple transcripts that differ between them. These data lay a foundation for further sub-classification and integration with rapidly evolving understanding of functional distinctions between reticulospinal subtypes (Ruder et al., 2021; Usseglio et al., 2020). Overall, across the supraspinal connectome these new data enhance the collective understanding of neural diversity by linking the transcriptional state of cells to the location of their cell bodies, and by extension to previously established understanding of physiological roles.

### Implications for injury and diseases

Damage to the spinal cord disrupts many ascending and descending tracts that carry a wide range of motor, sensory, and autonomic functions. Our recent meta-analysis indicated that research has historically focused disproportionately on major motor pathways such as the corticospinal, rubrospinal, and reticulospinal(Blackmore et al., 2021). Strong interest in main pathways is certainly warranted, yet arguably has led to a more limited understanding of many additional supraspinal pathways that carry functions also of value to individuals with spinal injury. As scRNA-seq opens new opportunities distinguish different types of neurons, it is increasingly practical to study a broader set of cell types impacted by spinal injury, in alignment with the needs of individuals with spinal injury. In this context, the current data serve as a needed foundation for at least three research directions.

First, identifying marker transcripts for selected populations is an initial step in creating cell-type specific vectors for gene manipulation. In principle, by identifying transcripts expressed in only in one supraspinal cell type but excluded from all others, it is possible to identify the relevant promoter or enhancer sequences and incorporate them into vectors to recapitulate selective gene expression. This strategy has been used for selective gene expression in layer V cortical neurons and specific inhibitory subclasses (Graybuck et al., 2021; Vormstein-Schneider et al., 2020). Importantly, as we have emphasized throughout, the markers described here are specific within the context of the supraspinal connectome but can be found elsewhere in the brain in non-supraspinal neurons. Thus, the preferred strategy would be to design vectors with cell-type specific enhancer elements and then deliver them to the spinal cord in retrograde fashion to avoid this off target expression. Delivering optogenetic, chemogenetic, or therapeutic transgenes in this way offers a strategy to parse discrete functions or selectively stimulate repair in selected supraspinal populations.

Second, the current data provide baseline information for future efforts to characterize transcriptional changes triggered by axotomy or disease. It has been known for decades that different supraspinal cell types display highly distinct regenerative responses after spinal injury and after treatment, for example seminal work that showed regeneration of brainstem-spinal axons, but not corticospinal, into peripheral nerves grafted to the spinal cord (David and Aguayo, 1981; Richardson et al., 1984) One possibility is that underlying these different growth responses are variations in the transcriptional response to injury. The present data provide an initial classification of transcriptionally distinct supraspinal cell types and, critically, a population-by-population baseline to identify transcriptional changes after damage or disease.

Finally, the present data can be mined to generate hypotheses regarding functional differences between supraspinal populations. As one example we have examined receptors for axonal growth and guidance cues, both endogenous and exogenous. We find that supraspinal populations generally express receptors for neurotrophins BDNF and NT-3 but not NGF, consistent with an extensive literature regarding the growth effects of these ligands after spinal injury (reviewed in (Keefe et al., 2017). Supraspinal populations also generally lacked receptors for GDNF, which may explain findings that it stimulates axon growth from propriospinal but not supraspinal axons (Anderson et al., 2018; Deng et al., 2013). Interestingly, we also find that most supralumbar populations express Crim1, a transmembrane domain protein that contributes to lumbar targeting in CST neurons (Sahni et al., 2021a), hinting at a wider role in directing axons from a variety of cell types to lumbar targets.

In contrast, some receptors were highly variable between supraspinal populations, hinting at differential responses to cues. For example, semaphorins are inhibitory cues expressed in injured spinal tissue (Shim et al., 2012; Ueno et al., 2020). CST neurons express high levels of semaphorin receptors Nrp1and Plxna2 and interfering with semaphorin / Plxna2 signaling enhances post-injury sprouting and reduces axon retraction in CST neurons (Shim et al., 2012; Ueno et al., 2020). We find, however, that most other supraspinal populations expressed very low levels of semaphorin receptors, indicating that this inhibition may be specific to CST axons; other supraspinal tracts may benefit less after relief from semaphoring inhibition. Netrin-1, which can attract axons via the Dcc receptor or to repulse axons via receptors in the Unc5 family, is expressed by spinal oligodendrocytes after injury and has been studied both as an endogenous inhibitor of axon growth (Löw et al., 2008) and a potential reparative factor. Interestingly, compared to other populations CST neurons stood out as expressing very low levels of Dcc and higher levels of Unc5a, hinting at a more negative response to this cue. As mentioned above, these predictions are subject to important caveats regarding the tenuous link between mRNA abundance and protein expression and/or localization. Nevertheless, they illustrate the broader point that careful examination of transcriptional differences between supraspinal populations can be used to guide future experiments to test differential function.

Other important caveats apply to these data. First, we were constrained by technical considerations to exclude some populations that were too small or too dispersed for rapid collection and sorting. Notable examples include the pontine reticular formation and a small population in the dorsal medulla in the vicinity of the solitary nucleus. A second caveat regards the tropism of AAV2-retro. The lumbar spinal cord receives substantial descending input from serotonergic, noradrenergic, and gabaergic populations, yet together they comprised less than three percent of cells retrogradely transduced with AAV2-retro. The relative inefficiency of AAV2-retro in serotonergic and noradrenergic cell types has been noted previously by our lab and others (Ganley et al., 2021; Tervo et al., 2016; Wang et al., 2018). The current data thus add to accumulating evidence that the tropism of AAV2-Retro favors glutamatergic neurons and points toward the importance of developing additional retrograde vectors with tropism appropriate for additional cell types. Overall, to further expand understanding of the supraspinal transcriptome it will be necessary to optimize sample preparation to capture residual populations and to deploy labeling strategies with broader tropism. Nevertheless, the present data offer a significant advance in information regarding into the transcriptional state of a wide range of neuronal populations that communicate with the lumbar spinal cord and offer new insights into potential physiological differences.

## Acknowledgments

This work was supported by grants from NINDS and the Bryon Riesch Paralysis Foundation, The Miami Project to Cure Paralysis, and the Buoniconti Fund. We thank Angela Schmoldt (UW-Milwaukee) for bio-analyzer assistance and the UW-Madison Biotechnology Center for sequencing. We also thank James Choi, Dr. Dmitry Velmeshev, Dr. Kevin Park, and Dr. Jae Lee for insight and critical evaluation of the work.

