## Supplementary figures and images for "Single nuclei analyses reveal transcriptional profiles and marker genes for diverse supraspinal populations"

### Extended Data 7-1

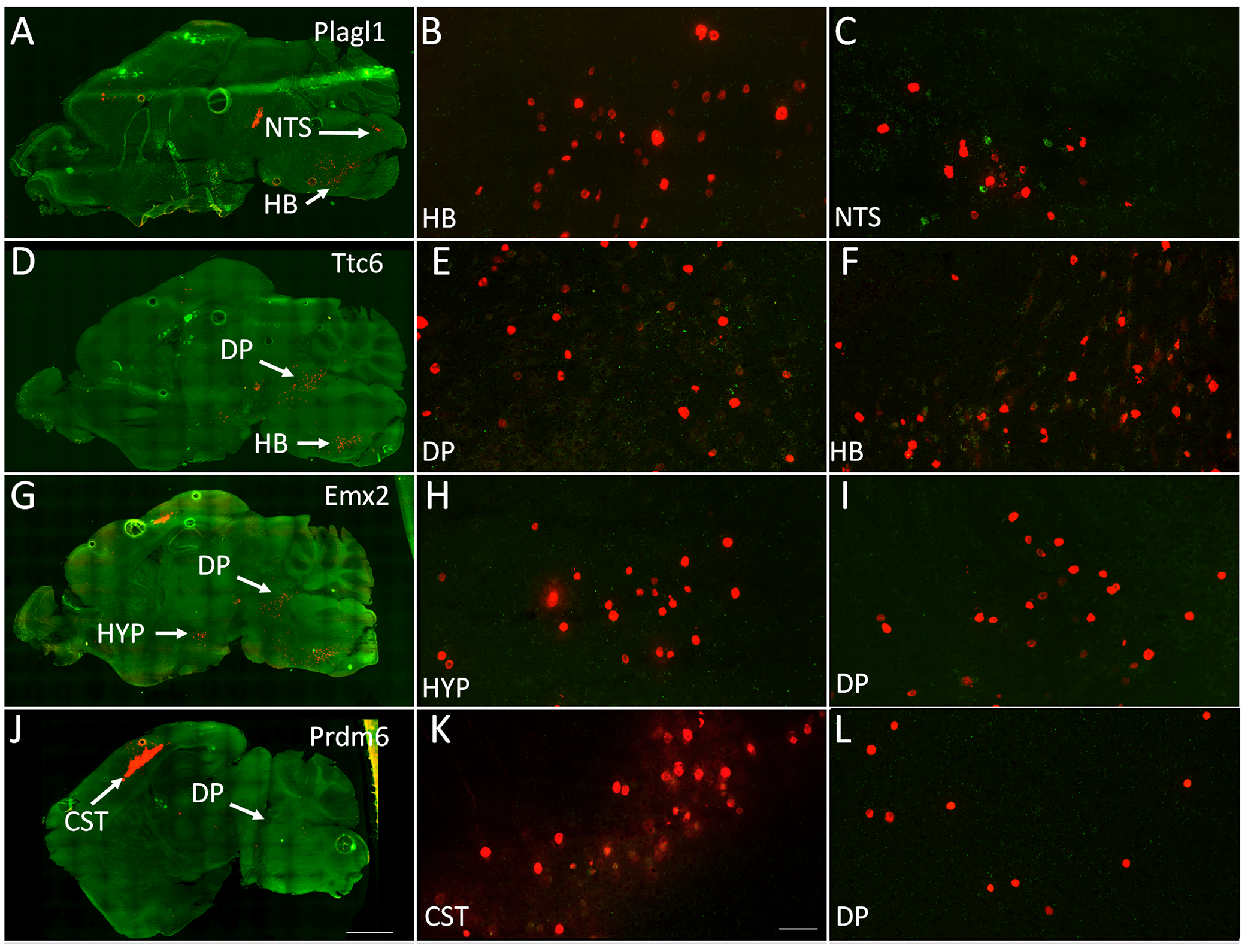

### Extended Data 8-1

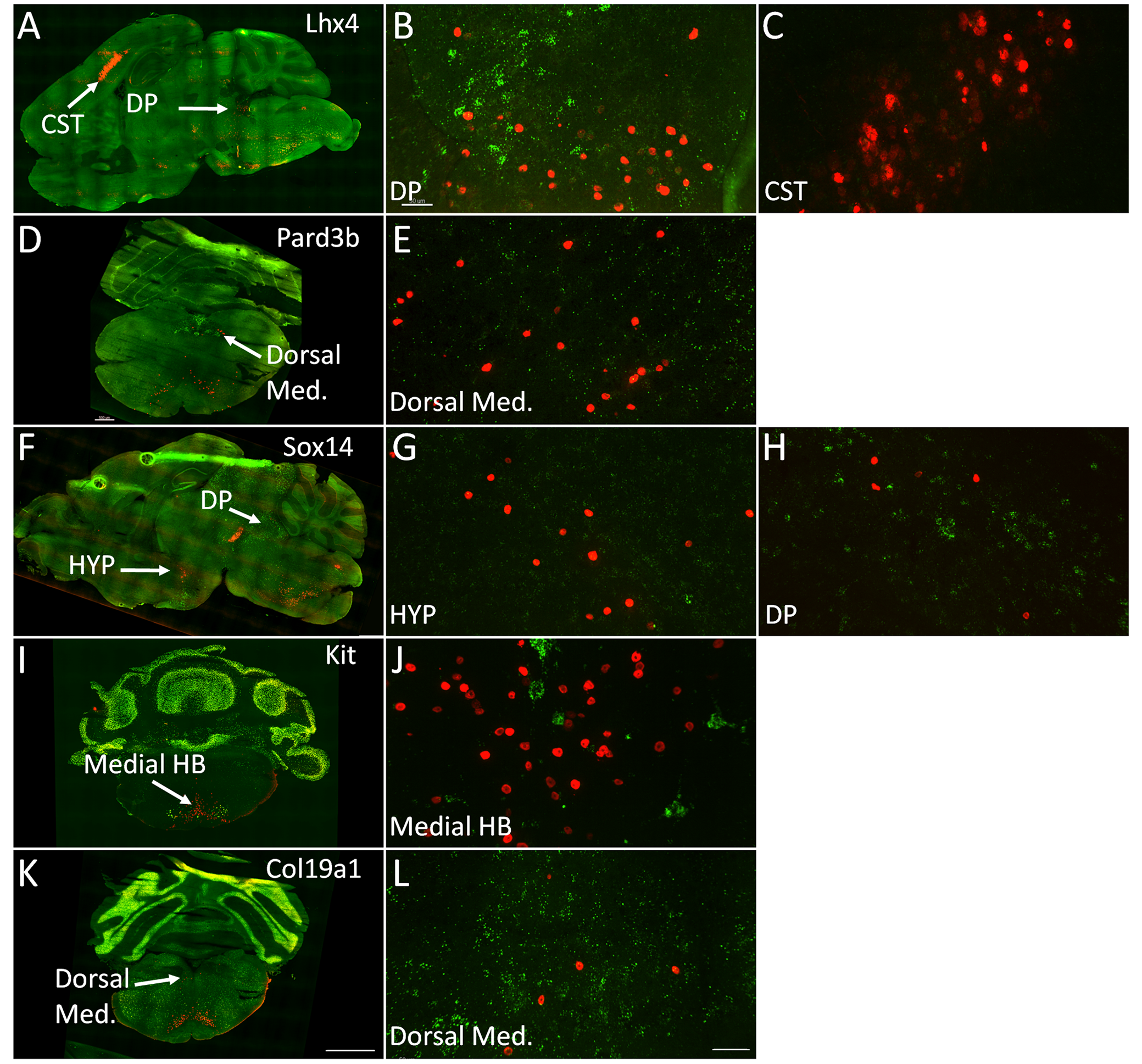
